# Humidity as a non-pharmaceutical intervention for influenza A

**DOI:** 10.1101/273870

**Authors:** Jennifer M. Reiman, Biswadeep Das, Gregory M. Sindberg, Mark D. Urban, Madeleine E.M. Hammerlund, Han B. Lee, Katie M. Spring, Jamie Lyman-Gingerich, Alex R. Generous, Tyler H. Koep, Kevin Ewing, Phil Lilja, Felicity T. Enders, Stephen C. Ekker, W. Charles Huskins, Hind J. Fadel, Chris Pierret

## Abstract

Influenza is a global problem infecting 5-10 % of adults and 20-30 % of children annually. Non-pharmaceutical interventions (NPIs) are attractive approaches to complement vaccination in the prevention and reduction of influenza. Strong cyclical reduction of absolute humidity has been associated with influenza outbreaks in temperate climates. This study tested the hypothesis that raising absolute humidity above seasonal lows would impact influenza virus survival and transmission in a key source of influenza distribution, a community school. Air samples and objects handled by students (e.g. blocks and markers) were collected from preschool classrooms. All samples were processed and PCR used to determine the presence of influenza and its amount. Additionally samples were tested for their ability to infect cells in cultures. Deliberate classroom humidification (with commercial steam humidifiers) resulted in a significant reduction of the total number of influenza positive samples (air and fomite), viral copy number, and efficiency of viral infectivity. This is the first prospective study suggesting that exogenous humidification could serve as a scalable NPI for influenza or other viral outbreaks.

**Author summary:** Human influenza infections have a substantial impact on society (including lost productivity and medical costs). Children, 3-4 years of age are the main introducers and spreaders of influenza within a household and community. There is evidence from laboratory and epidemiological studies that suggests that low humidity in winter (in temperate climates) may increase the ability of influenza virus to survive and spread between individuals. We wanted to know if added in humidity (through steam humidifiers) could reduce the amount of influenza present and its spread within preschool classrooms (students aged 3-4 years)? Additionally, we looked at the infectivity of the influenza isolated and if there were differences in the number of students with influenza-like illnesses during our study. We show that humidification can reduce the amount of influenza present within samples from preschool classrooms and that there were fewer infectious samples compared to non-humidified rooms. There were small numbers of students ill with influenza like illnesses during our study so additional studies will need to look further at humidification as a way to reduce influenza infection and transmission.

## Introduction

Non-pharmaceutical interventions (NPIs) can complement traditional vaccination and anti-viral medications for infectious disease. NPIs are particularly significant for diseases like influenza, which have not shown to be well managed by vaccination alone. Worldwide, annual influenza epidemics are estimated to infect 5-10 % of adults and 20-30 % of children resulting in 3 to 5 million cases of severe illness and 250,000 to 500,000 deaths[1]. These infections account for 10% of global respiratory hospitalizations in children under 18 years of age[2]. Direct medical costs (2015) for influenza for inpatient care have been estimated at $3,650-$9,660 per case (reviewed in[3]) and a total economic burden of influenza exceeding $87 billion per year in the USA[4]. In Minnesota, USA, the 2014-15 influenza season was especially severe with 4,202 people hospitalized with laboratory-confirmed influenza infection and 10 confirmed pediatric influenza-related deaths. This was the highest rate reported for the past six seasons including the H1N1 influenza pandemic in 2009[5]. Influenza and respiratory syncytial virus (RSV) accounted for over 50 % of hospitalizations (all ages) from respiratory infections with an additional 706 outbreaks of influenza-like illness (ILI) reported in schools[5]. This increase in influenza was attributed to antigenic drift, leaving the vaccine largely ineffective. The 2014-15 influenza season illustrates why alternative approaches such as NPIs could prove to be valuable in disease prevention.

The probability for influenza transmission increases in situations where many people are in enclosed spaces and in close proximity such as airplanes and naval ships[6]. Children play a critical role in the transmission of acute respiratory infections within a community[7]. Survival and transmission of influenza may be impacted by the droplet size of airborne influenza. Larger droplets settle out of air at a more rapid rate, and do not penetrate as deep into the respiratory tract to seed infection[8]. Under laboratory conditions, others have shown that infectious influenza virus can persist from hours to days (17 days on banknotes with respiratory secretions[9]) in objects as varied as surface dust[10], cloth, pillow case, soft fabric toy, light switch material, formica, vinyl, and stainless steel[11, 12]. Researchers have demonstrated transfer of infectious influenza A from stainless steel surfaces (up to 24 hours) or from paper tissues (15 minutes) to hands, while remaining infectious on hands for 5[13] - 60[12] minutes.

The mortality and transmission rates of influenza A have been associated with decreased absolute humidity (AH) [14]. This epidemiological correlation suggests that deliberate increases in AH could be a potential NPI to reduce the spread of influenza and other viruses. One approach is to maintain relative humidity (RH) between 40-60 %, the proposed optimal range for reducing growth opportunities for viruses, bacteria, and fungi[15]. Our previous study, demonstrated that classroom humidification to RH of 40-60 % may be a feasible approach to increase indoor AH to levels with the potential to reduce influenza virus survival (a target of 10mb) and transmission as predicted by modeling analyses[16].

Community schools are promising locations for potential use of humidity as a NPI due to the role of children as a key source of influenza transmission from the community into a household[4]. We developed a novel in-school sampling process (S1 Fig) sensitive enough to detect influenza presence, quantity (viral RNA), and infectivity.

Outside of the laboratory, no prior studies have tested the potential for humidification to serve as an NPI. A few groups have succeeded in collecting influenza (RNA) from air samples in health care settings[17, 18]. One group also detected influenza (RNA) within air samples at a day-care facility (babies’ and toddlers’ rooms) and on board airplanes across the USA[18]. However, these studies did not detect and isolate infectious influenza from fomites in field conditions. This study investigated the presence and infectivity of influenza A in active preschool classrooms under control and humidified conditions.

## Results

At the end of the 2015-2016 influenza season, an analysis of MN hospitalizations attributed to influenza (data supplied by MN Department of Health via FluSurv-NET)[19] revealed three troughs in atmospheric absolute humidity with the largest one (February 14, 2016)[20] preceding the peak of the seasonal influenza outbreak (Fig. 1A). The peak for confirmed hospitalized influenza cases (confirmed positive) was the week ending March 12, 2016.

**Fig. 1.**
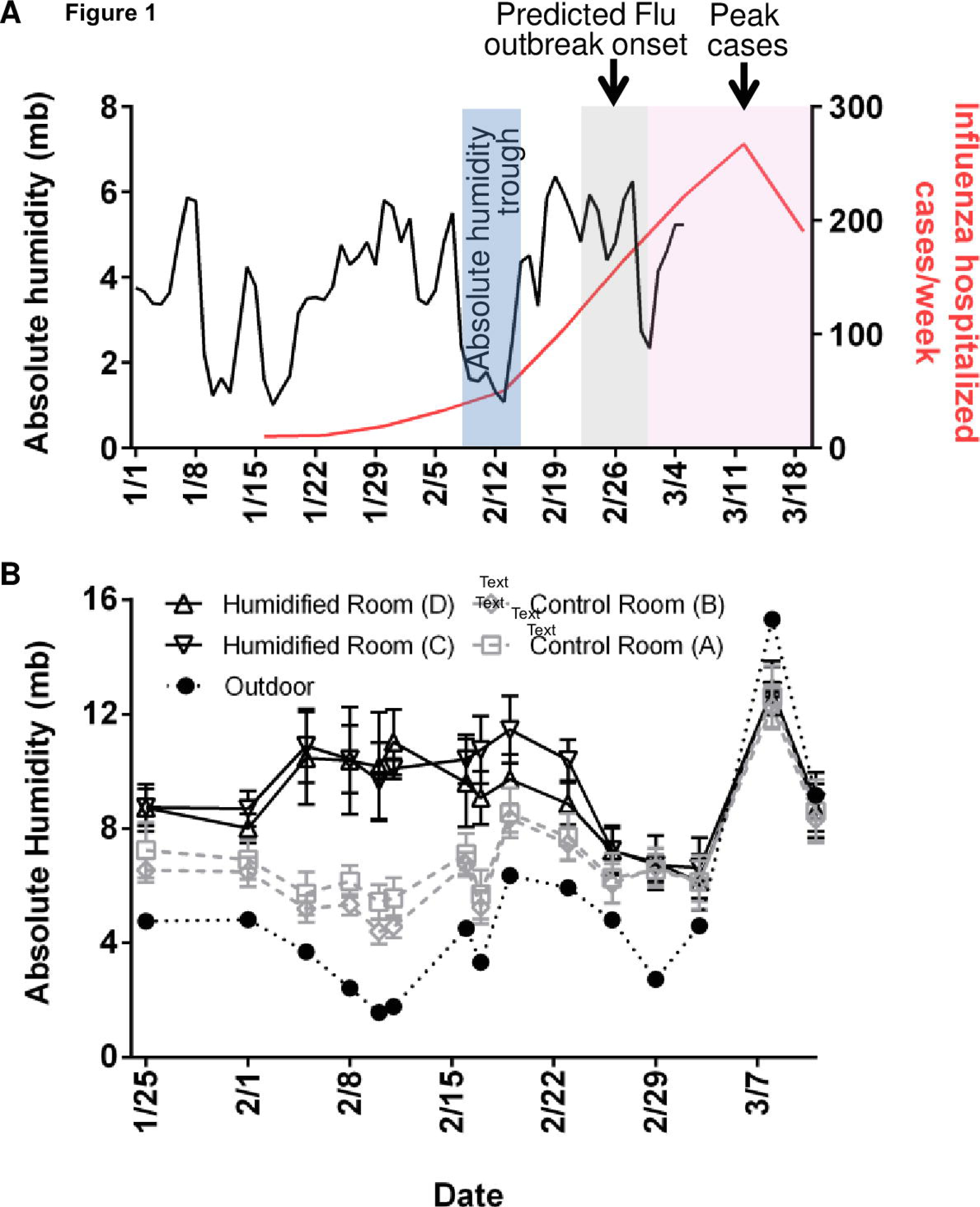
Absolute Humidity and influenza hospitalized cases. (**A**) Outdoor absolute humidity (AH) values (n=65; one measurement per day) from Rochester, MN and influenza hospitalized cases in MN (n=1070, week ending in January 16th-March 19th). Applying the national trend model described by Shaman et al.[31] to the local humidity and illnesses, onset of influenza followed the predicted delay of 10-16 days (grey box) after an absolute humidity trough (blue box). Peak cases follow (pink box), as there is an incubation period of 1-4 days with viral shedding up to 7 days after symptoms resolve. (**B**) AH in 4 preschool classrooms (average of two sensors from 10 minute intervals over 150 minutes (n=16 per sensor, room D) or (n=17 per sensor, rooms A,B, C) per class period per sensor). Center values are mean of both sensors during class time and error bars are s.d. and corresponding outdoor AH (n=15, 1 per day) on the 15 days of sample collection. Humidifiers were running in humidified rooms through sample collection on February 23.

Elevated classroom humidification was maintained at an average of 9.89 mb in humidified rooms compared to 6.33 mb in control rooms (January 25 through February 23). AH was targeted near 10 mb based on previously demonstrated achievable levels in classrooms and a calculated 1-hour virus survival of 35 % (down from 75 % when AH ∼3-4 mb). Samples positive for influenza A virus are depicted in Table 1.

**Table 1.**
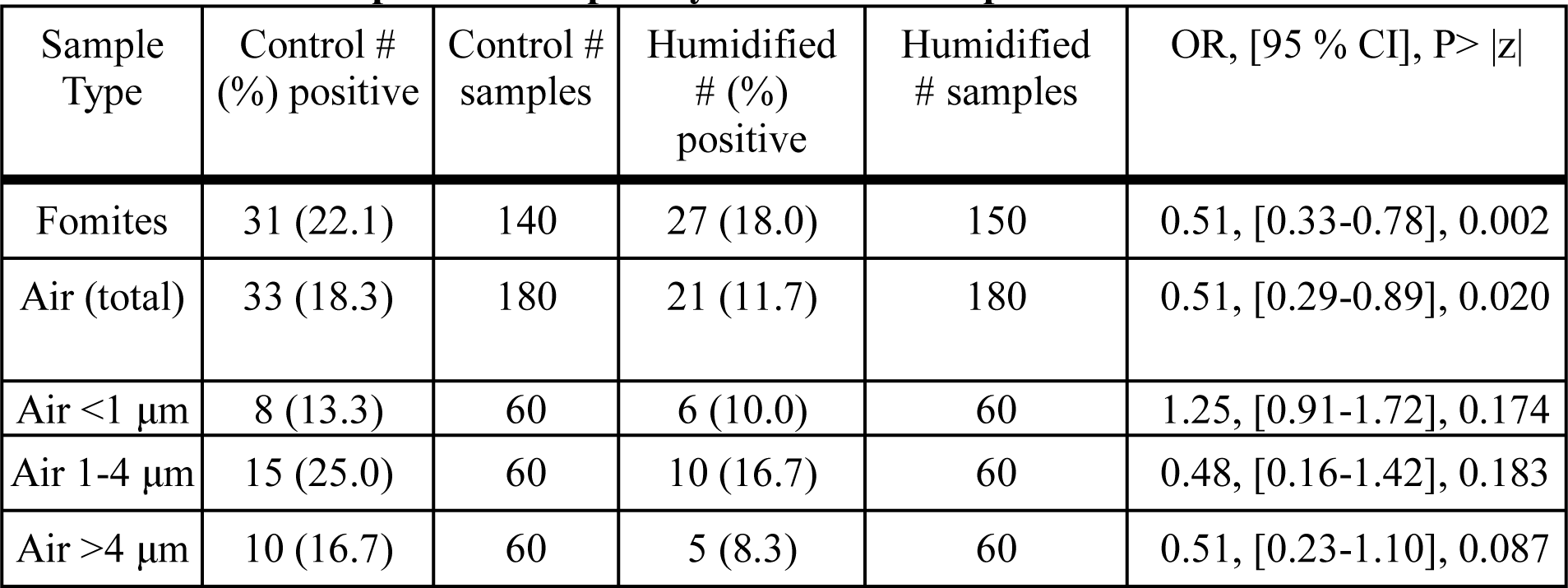
**Influenza A positive samples by RT-PCR from preschool.**

The number of positive samples in each sample type is indicated, the total number of samples collected and the % positive (in parentheses) for control and humidified rooms. Statistical analyses of OR, 95 % CI and P>|z| are indicated.

A total of 650 samples were collected (320 in control rooms, 330 in humidified rooms) of which 112 (17 %) were positive for influenza A virus by RT-PCR (S2A-2B Figs). There were fewer influenza A positive samples in humidified rooms compared to control rooms for both fomites and for total air. However, when individual sizes of air particles were examined, differences did not achieve statistical significance. The distribution of influenza A positive samples within the different sizes of air particles varied with the greatest percentage of positive samples within the 1-4 μm size for both control and humidified rooms (Table 1).

Quantitative RT-PCR of influenza A virus for copy number revealed a significant reduction in mean copy number in humidified rooms compared to controls for fomites (P<0.001) and air (total) (P<0.001). Review of qRT-PCR data at individual particle sizes revealed that air <1 μm (P=0.010), and air > 4 μm (P=0.011) experienced a significant reduction (Fig. 2A and Table S1). Mean copy number for air between 1-4 μm (P=0.208) trended lower in humidified rooms but was not statistically different from control rooms at a 95 % CI. Viral copy number varied among different sample types with fomites having the highest copy number followed by air 1-4 μm particles (Table S1).

**Fig. 2.**
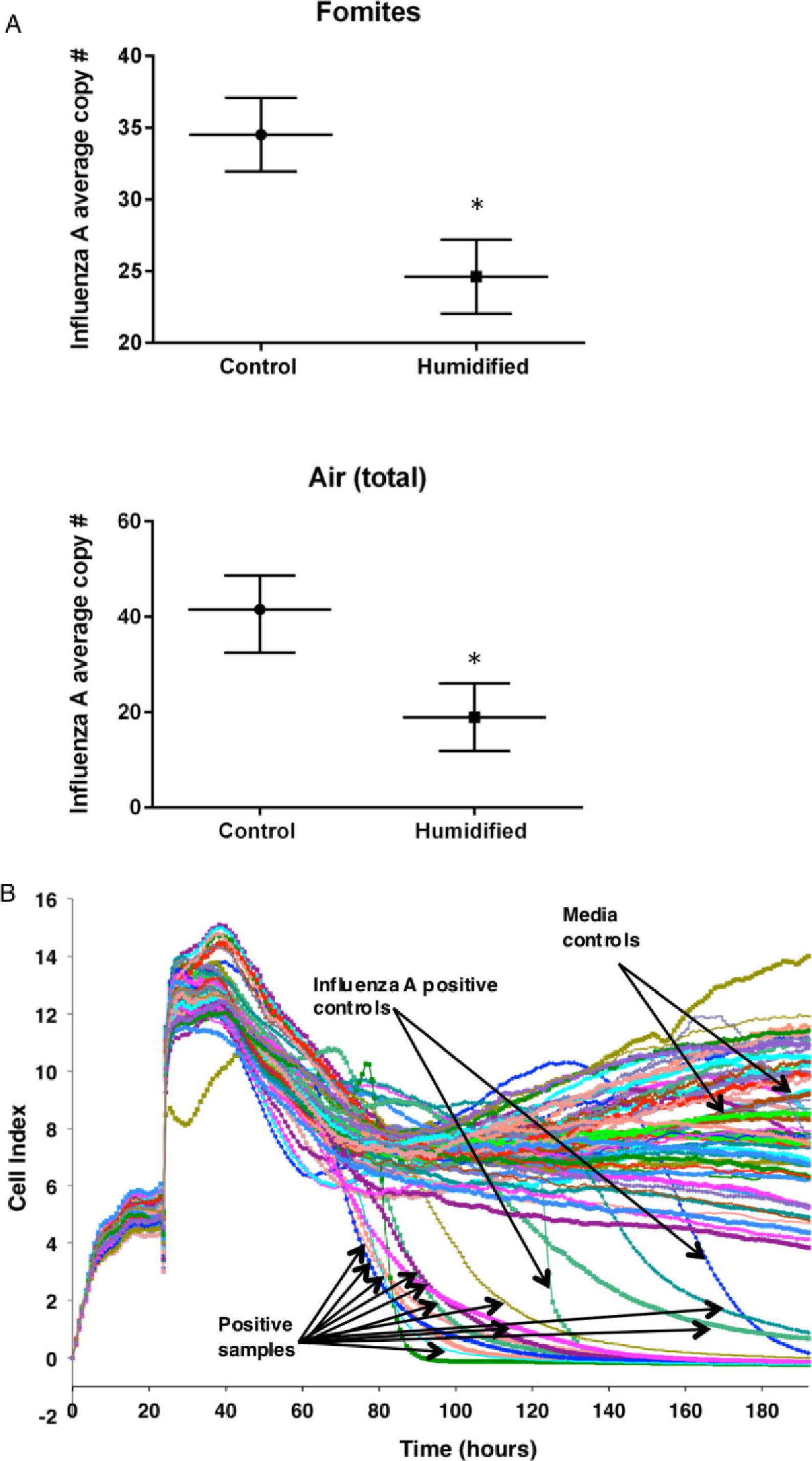
Influenza A (NS1) average copy number and infectivity of positive samples in electrical impedance assay. (**A**) Horizontal bars indicate mean copy number and error bars are 95% CI. Fomites control, n= 31; Fomites humidified, n=27; Air (total) control, n=33; Air (total) humidified, n=21. (**B**) Each line indicates a sample (well) for infectivity. MDCK cells were added at time 0 hours and media changed to samples (n=27 in triplicate), influenza A positive control dilutions (6) in duplicate or control media in triplicate at 24 hours. Cell indices that returned to 0 indicated cell death (infectious). Arrows indicate individual samples. * indicates P<0.001.

Electrical impedance assay [21] revealed 30 % of RT-PCR positive classroom samples to be infectious (Fig. 2B and Fig. S3). There were a smaller percentage of infectious samples from the humidified rooms (19 %) than the control rooms (81 %). The corresponding average viral copy number in the infectious samples in the humidified rooms was 30 and 40 in control rooms.

The majority of air particles were less than 1μm size for both humidified and control rooms. Average particle numbers decreased with increasing size in both sets of rooms (<1 μm: 45500 control vs 70400 humidified), (1-4 μm: 2820 control vs 4060 humidified), (>4 μm: 139 control vs 227 humidified). Particles generated from the humidifier and the average counts from 7 independent measurements over a 22-minute period showed more than 96% of the particles provided by humidification were less than 1 μm. Yet, humidified rooms showed a near doubling of both 1-4 μm, and >4 μm air particles. This indicates a likely combination of the small particles added by humidification with present (and potentially influenza-bearing) particles. Larger particles remain airborne for less time (4 μm takes 33 minutes to settle 1 meter in still air versus 1 μm takes 8 hours)[17] and are unable to reach as deep within airways so are thought to be less pathogenic[8].

From January 25-March 11, 2016, 10 influenza-like illnesses (ILI) from absent students were recorded by school personnel. Seven of these absences were from control (non-humidified) rooms and three were from humidified rooms. As shown in Table 2, in a per-room comparison of humidified and control rooms the % positive (# PCR+ /total samples per room), % infective (# positive by infectivity/ # PCR positive tested per room), and % of students with ILI absences (# students with fever + cough or fever + sore throat / 32 days of classes) all were lower in humidified rooms than control rooms.

**Table 2.** **Influenza A positive and infectious samples and ILI absences by room.**

|  | Control |  | Humidified |  |
| --- | --- | --- | --- | --- |
| Influenza A | Room A | Room B | Room C | Room D |
| % positive<br>(# PCR+ / total<br>samples per room) | 33/160=<br>21 % | 31/160=<br>19 % | 27/165=<br>16 % | 21/165=<br>13 % |
| % infective<br>(# positive by<br>infectivity/ # positive<br>tested per room) | 6/15=<br>40 % | 7/12=<br>58 % | 2/7=<br>29 % | 1/11=<br>9 % |
| % students ILI*<br>(# students ILI/<br>32 days of classes) | 3/32=<br>9 % | 4/32=<br>13 % | 2/32=<br>6 % | 1/32=<br>3 % |

Percentage of student absenteeism showed a logical flow from initial positive sampling. % positive is number PCR+ divided by total samples per room. % infective is number positive by electrical impedance assay divided by number PCR positive samples tested in electrical impedance assay. ILI indicates influenza-like illness absence based on symptoms reported. % of students with ILI absences is number of students out ill with fever plus cough at any point during the 32 days of classes (January 25-March 11, 2016) divided by the 32 classes. *Only absences within the timeframe of sample collection (January 25-March 11, 2016) are included.

## Discussion

This study monitored transmission of influenza A in preschool classrooms during the dry winter months (low indoor humidity), which correspond with peak respiratory virus infections in Minnesota. An increase in average AH from 6.33 mb in control rooms to 9.89 mb in humidified rooms (RH ∼42-45 %) was associated with a significant decrease in influenza A virus presence in fomite and air samples in humidified rooms compared to control rooms. Additionally, PCR-positive samples from humidified rooms exhibited lower infectivity than samples from control rooms. The decrease in infectivity could be driven by a lower viral load of the PCR-positive humidified room samples as compared to PCR-positive control room samples. This correlates with the lower copy number of influenza A PCR positive samples observed in humidified rooms. Enveloped viruses such as influenza are less stable in the environment than non-enveloped viruses and are more sensitive to higher relative humidity[22-24]. The exact mechanism underlying the action of humidity on the survival of the influenza virus is still not fully understood[25]. Several studies have hypothesized that surface inactivation for viruses with structural lipids may be due to denaturing of the lipoproteins found in enveloped viruses and phase changes in the phospholipid bilayer leading to cross-linking of associated proteins[22]. Additionally, relative humidity impacts the salt concentration within droplets and at low relative humidity (<50 %), the solutes crystalize and influenza viability was maintained[26].

Prior studies have largely focused on the influence of humidity (RH or AH) on influenza virus survival under laboratory conditions ([8, 24-27]). One study modeled influenza virus survival across varied ranges of ambient indoor AH and humidification levels achievable in school environments[16]. Based on these findings, we hypothesized that classroom humidification might be a feasible approach to increase indoor AH to levels that could decrease influenza virus survival and transmission. Taking classroom humidification the next step further we included collection of samples from classroom environments and demonstrated that humidified rooms exhibited fewer influenza A-positive samples and reduced copy number. Additionally, influenza A-positive samples were less infectious in humidified rooms. Together, these outcomes strongly support the hypothesis that deliberate humidification can mitigate influenza activity in a school environment.

In this study, humidifiers significantly increased the overall amount of particles, especially by increasing number of small air particles, finally leading to an increase in larger size particles, thereby rendering a shift in distribution towards largest particles, >4 μm. This could be attributed to the aggregation of particles in the humidified air, reducing the time of air suspension and transmission via inhalation. These findings enable a better understanding of the impact of humidity on influenza A virus survival (surfaces and objects) as well as virus transmissibility via aerosols by measuring the size of aerosol particles and the distribution of viruses within different sizes of particles.

Adding direct measurements of influenza (or other respiratory viruses) in a larger experimental population may reveal higher resolution outcomes to this approach. Furthermore, the impact of exogenous humidification on other viruses will need to be similarly measured to more fully understand the full potential impact of this NPI on other sources of communicable respiratory infections.

## Materials and Methods

### Study site and absence data

The study was conducted at Aldrich Memorial Nursery School, Rochester, MN (a preschool with students aged 2-5 years) from January 2016-March 2016. Classrooms of identical design (see S4 Fig) each with their own HVAC system for air handling were utilized. This study received prior IRB review and has been approved as an IRB #15-000476, an exempt study (Not Human Subject Research). Aldrich staff collected information on student absences (January 4, 2016-March 31, 2016) including ILI symptoms from students who were ill. ILI was defined as having fever plus cough or sore throat.

### Humidifiers, humidity measurements, and absolute humidity calculations

Model XTR (XTR003E1M) electrode steam humidifier with steam blower (SDU-003E) (DriSteem, Eden Prairie, MN) was installed in two experimental classrooms. The boiling of the softened tap water source provided decontamination of the steam distributed through the steam blowers. Classroom temperature and relative humidity were recorded every 10 minutes during the duration of the study with HOBO external data loggers, Model #U12-012 (Onset Computer Corporation, Bourne, MA). Two data loggers were installed per classroom, placed on the interior walls and set on top of the bulletin boards at a height of 2.032 meters. Data exported to Excel using HOBOware software (Onset Computer Corporation, Bourne, MA). Outdoor temperature and relative humidity was obtained from the North American Land Data Assimilation System (NLDAS) project[20]. Absolute humidity was calculated using Excel software using formulas as previously described[16].

### Bioaerosol sampler

NIOSH two-stage bioaerosol cyclone samplers[28, 29] collected air samples and separated them into three size fractions (>4 μm, 1-4 μm, and <1 μm) at a flow rate of 3.5 L / minute. NIOSH samplers were connected to AirChek XR5000 personal air sampling pumps, Model 224-PCXR4 (SKC Inc., Eighty Four, PA). Flow rate was calibrated using a Mass Flowmeter 4140 (TSI, Shoreview, MN), prior to each run of 150 minutes. Falcon conical tubes (15mL) (Corning, Corning, NY), 1.5 mL Fisherbrand microcentrifuge (Fisher Scientific, Pittsburgh, PA) and 37 mm hydrophobic Fluoropore PTFE membrane with a 3.0 μm pore size (EMD Millipore, Billerica, MA) were used to collect 3 different size fractions. Air sampling pumps were placed inside plastic ammunition boxes lined with mattress topper material (donated by Rest Assured Mattress Co, Rochester, MN). Cyclone samplers were affixed to the outside box surface with Industrial Strength Tape Strips (Velcro, Manchester, NH).

### Particle Counts

After class dismissal, particle counts were measured in the center of each classroom at a height of 91 cm for 1 minute using a Six Channel Handheld Particle Counter, Model 23v750 (Grainger, Lake Forest, IL). Sizes of particles measured were 0.3, 0.5, 1, 2.5, 5, and 10 μm. Particle size data was binned into sizes to match those collected by the NIOSH samplers such that <1 μm included 0.3 and 0.5 μm sizes, 1-4 μm included 1 and 2.5 μm and >4μm included 5 and 10 μm. Particle counts were also measured before and after humidifier turned on in a pilot test.

### Air samples

NIOSH samplers were dissembled in a BSLII biosafety cabinet. Collection tubes received 1 mL of infection media. Filters were retrieved from inside black polypropylene filter cassettes opened with a Stainless Steel SureSeal Cassette Opener (SKC Inc., Eighty Four, PA) and placed inside a 15 mL conical tube containing 1 mL infection media. The air filter was pushed down into the media using a sterile 1 mL serological pipette. Samples were placed on ice until ready for further processing.

### Fomites

25 % cotton linen paper (Southworth, Neenah, WI) wrapped objects were provided to students. Objects included markers and a variety of wooden toys including blocks, rolling pins patterned wheel press and stamping cubes (Melissa & Doug, Wilton, CT). Additional items present in the classroom were also wrapped including rolling pins, hard rubber brayers, and plastic pizza cutters. After play, paper was transported back to BSL2 laboratory.

### Processing of fomites and air samples

Papers from classroom objects were lightly dusted with fingerprinting powder (Hi-Fi Volcano Latent Print Powder, Sirchie Youngsville, NC). Once fingerprint was identified, a portion of paper (∼3.5-4 cm^2)^ was removed and placed into a 15 mL conical tube containing 1 mL of infection media. Samples were placed on ice until ready for further processing.

Both fomites (paper) and air samples (in media) were vortexed briefly and incubated on ice for 15 minutes. Samples were vortexed again prior to centrifugation for 20 minutes at 4 °C at 3313g (15 mL tubes) and centrifuged at room temperature at 1520g in a microcentrifuge (1.5 mL tubes). Liquid was removed from 15 mL conical tubes and transferred to 1.5 mL conical tubes. The supernatant from these samples was frozen at −80 °C and used for subsequent RNA isolation and qRT-PCR.

### Influenza A virus controls and infectivity assay

Viral stocks of influenza A (H3N2) were provided by the Clinical Virology Laboratory, Mayo Clinic, Rochester, MN. Samples were thawed and used to infect bulk cultures of MDCK. Influenza A infections followed published methods[30], except DMEM media used instead of MEM. The contents of the flask were collected once cells became non-adherent and centrifuged 4 °C at 3313g for 10 minutes to pellet cellular debris. Aliquots of supernatant were stored at −80°C. Samples were assayed for influenza A virus infectivity by electrical impedance assay[21] run on an xCELLigence RTCA MP instrument (ACEA Biosciences, Inc., Sand Diego, CA). After calibration with media, wells were seeded with MDCK cells. Twenty-four hours later, media was removed and the cells were washed with PBS. Wells were inoculated with serially diluted influenza A virus (positive controls), media (negative controls) and samples (tests). Readings were taken every 15 min for 7 days post inoculation. Using influenza A virus stock, dose dependent decline in cell index was demonstrated with 1:1000 dilutions, and gradual decline at 1:10000 and 1:100000 dilutions, whereas further dilutions did not impact cell index any differently than media controls (Fig. 2B and Fig. S3).

### Viral RNA isolation and detection using quantitative reverse transcription PCR (qRT-PCR)

The supernatant from fomite and air samples was thawed and 140 μL used for viral RNA isolation with the Viral RNA isolation kit (Qiagen) as per the manufacturer’s instructions, followed by RT-PCR analyses. The nonstructural (NS1 gene) sequence of influenza A was used for virus detection. See S2 Table for list of primers, product sizes and annealing temperature information. See S5 Figure for detection of viral RNA in multiplexed qRT-PCR. Briefly, SYBR green qRT-PCR was performed as described by the manufacturer (Qiagen SYBR green Quantitect kit). The PCR thermal profile consisted of an initial cDNA step of 30 minutes at 50°C followed by 15 minutes at 95 °C and 30 cycles of 30 seconds at 95 °C, 30 seconds at 56.5 °C, and 30 seconds at 72 °C. Detection, quantification and data analysis were performed in the CFX manager real-time detection system (Bio-Rad).

### In-vitro transcription of influenza A viral RNA

Though Influenza A and B and RSV were included in this analysis, low numbers of positive samples for RSV and Influenza B moved the focus of the study to influenza A only. Quantification of influenza A RNA was performed using in vitro transcription (IVT). The NS1 gene was amplified from a H3N2 influenza A virus isolate from infected MDCK cells using primers containing T7 promoter sequence in the forward sites: Inf AF: 5′-ACTGCTTAATACGACTCACTATAGGGAGATTTCACCGAGGAGGGAGCA -3′, Inf AR: 5′-CCTCCGATGAGGACCCCAA -3′;. The amplified NS1 gene was in-vitro transcribed with T7 RNA polymerase (Megascript T3 kit, Ambion, ABI) according to the manufacturer’s instructions for synthesizing short transcripts (105 bp) with the following modifications: incubation time increased to 8 hours and the enzyme mixture and template RNA was increased by three-fold of its original concentration. The synthesized RNA pellet was suspended in 0.1% diethylpyrocarbonate-treated water. RNA transcripts were purified, quantified and mixed with nuclease free water for preparation of positive controls in the range from 10^1^-10^7^ copies for the standard curve development. The amount of IVT-generated fragments was determined using the NanoDrop ND2000 Spectrophotometer (NanoDrop Technologies, INc., Wilmington, DE) and converted to molecular copies according to the formula:

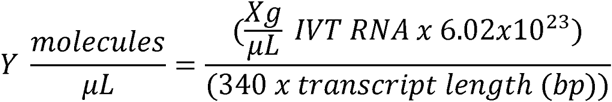

The detection limit of the SYBR green real time RT-PCR assay was determined by testing serial ten-fold dilutions of the in vitro transcribed influenza A viral RNA ranging from 10^1^ to 10^7^ copies/µL. Cycle-threshold (Ct) values were plotted against the RNA copy number to construct the standard curve. The viral copy numbers of the processed samples were estimated by plotting the respective Ct values on the standard curve. See S6 Figure for standard curve example.

### Detection and quantification

Using serially diluted in vitro transcribed influenza A RNA, viral particles were detectable ranging from 10^1^ - 10^7^ RNA copies using the described assay. qRT-PCR sampling was sufficiently sensitive to detect as few as 7 RNA copies.

## Statistical analyses

Percentage of samples positive for influenza A (Table 1): To account for within-room clustering for class cohorts, generalized estimating equations were utilized with a binomial family and logit link. Data are reported as odds ratio of positive test result for humidified rooms as compared to control rooms, with a 95% confidence interval and p-value. Odds ratios below one (with statistical significance) demonstrate protective effects of humidification in that positive influenza was less likely to be obtained by the relevant capture system. Capture systems for influenza included paper and air (total, >4 μm, 1-4 μm, <4 μm); results were treated as positive for any capture within the room to minimize the effect of the different number of collectors per room for the paper and air capture systems.

Mean copy number of influenza A positive samples (Table S1): To account for within-room clustering for class cohorts, generalized estimating equations were utilized with a gaussian family and identity link. Data are reported as the mean (standard deviation) of copy number for humidified rooms as compared to control rooms, with a 95% confidence interval and p-value. Capture systems for influenza included paper and air (total, >4 μm, 1-4 μm, <4 μm).

## Data Availability

The data that support the findings of this study are available from the corresponding author upon request.

## Contributions of authors

AG/GMS/HJF/KE/PL/WCH/SCE/CP designed the study. BD/JMR did the sample collection, all experiments, analyses of results, and drafted the manuscript. GMS provided analysis of indoor humidity data and procured air sampling equipment from NIOSH/CDC. GMS, KMS, MDU, and HJF assisted with cell culture infectivity assays. BD and MDU performed RNA isolation. MEMH directed UMR undergraduate work-study students in processing of paper wrapped objects. HBL assisted with air particle analyses. JLG analyzed student absence/attendance data. ARG conducted initial pilot study at Aldrich winter-spring 2015. THK provided outdoor humidity data and assisted with humidity analyses. KE provided the student data, turned on/off humidifiers and made adjustments to settings, and assisted with school management. PL provided humidifiers. SCE helped with analyses and trouble-shooting and revised the manuscript. WCH provided wording for collecting ILI school data and revised the manuscript. HJF provided BSL-2 laboratory oversight, and assisted in analyses. CP conceived the study, helped with study design, analyses and drafted the manuscript. FTE provided statistical analyses.

## Acknowledgements

The authors would like to thank the staff, students, parents and board of directors at Aldrich Memorial Nursery School for their support in allowing us to conduct this study. We acknowledge the support from DriSteem for supplying the humidifiers and covering the cost of humidifier installation for this study. We also recognize Chuck Dixon and Rest Assured Mattress Co, Rochester, MN for allowing us to try out different noise deadening materials and donating the material for our study. We thank Matt Binnicker, PhD and colleagues in the Clinical Virology Laboratory, Department of Laboratory Medicine and Pathology, Mayo Clinic, for providing Influenza A virus and other virus isolates and advice. Thanks to Bill Lindsley at NIOSH/CDC for supplying the NIOSH bioaerosol samplers for air sample collection. We also acknowledge the work of University of Minnesota-Rochester work-study students (Paige Diedrick, Ashlyn Stenberg, Ali Ali, Shidae Yang) as well as other InSciEd Out team members for wrapping of wooden blocks and markers with paper and cleaning up these objects after sample processing for future use. Thanks to Steve Offer and the laboratory of Robert Diasio, Mayo Clinic Cancer Center, for use of the xCELLigence MP equipment for the infectivity assay. The methodology flow chart figure was prepared by J. Mark Curry, Division of Creative Media, Mayo Clinic.

This work was supported by CTSA Grant Number UL1 TR000135 from the National Center for Advancing Translational Science (NCATS), a component of the National Institutes of Health (NIH). Its contents are solely the responsibility of the authors and do not necessarily represent the official views of the NIH. Additional funding was contributed by Mayo Clinic Pediatric and Adolescent Medicine [through WCH] and intramural grants from Mayo Clinic Infectious Disease [to HJF].

## References

1. WHO. Influenza (Seasonal) Fact Sheet N°211. 2014. [updated March 2014; cited 2015 October 15]. Available from: http://www.who.int/mediacentre/factsheets/fs211/en/.

2. Lafond KE, Nair H, Rasooly MH, Valente F, Booy R, Rahman M, et al. Global Role and Burden of Influenza in Pediatric Respiratory Hospitalizations, 1982-2012: A Systematic Analysis. PLoS Med. 2016;13(3):e1001977. doi: 10.1371/journal.pmed.1001977.

3. Ozawa S, Portnoy A, Getaneh H, Clark S, Knoll M, Bishai D, et al. Modeling The Economic Burden Of Adult Vaccine-Preventable Diseases In The United States. Health Aff (Millwood). 2016;35(11):2124–32. doi: 10.1377/hlthaff.2016.0462.

4. Molinari NA, IR Ortega-Sanchez, Messonnier ML, Thompson WW, Wortley PM, Weintraub E, et al. The annual impact of seasonal influenza in the US: measuring disease burden and costs. Vaccine. 2007;25(27):5086–96. doi: 10.1016/j.vaccine.2007.03.046.

5. Minnesota Department of Health Weekly Influenza & Respiratory Illness Activity Report- Summary of 2014-15 Influenza Season. 2015. [updated October 6, 2016; cited 2015 September 10]. Available from: http://www.health.state.mn.us/divs/idepc/diseases/flu/stats/2014summary.pdf.

6. Musher DM. How contagious are common respiratory tract infections? N Engl J Med. 2003;348(13):1256–66.

7. Brownstein JS, Mandl KD. Pediatric population size is associated with geographic patterns of acute respiratory infections among adults. Ann Emerg Med. 2008;52(1):63–8. doi: 10.1016/j.annemergmed.2008.02.009.

8. Tellier R. Aerosol transmission of influenza A virus: a review of new studies. J R Soc Interface. 2009;6 Suppl 6(Suppl 6):S783–90. doi: 10.1098/rsif.2009.0302.focus.

9. Thomas Y, Vogel G, Wunderli W, Suter P, Witschi M, Koch D, et al. Survival of influenza virus on banknotes. Appl Environ Microbiol. 2008;74(10):3002–7. doi: 10.1128/AEM.00076-08.

10. Edward DF. Resistance of influenza virus to drying and its demonstration on dust. The Lancet. 1941;238(6170):664–6.

11. Greatorex JS, Digard P, Curran MD, Moynihan R, Wensley H, Wreghitt T, et al. Survival of influenza A(H1N1) on materials found in households: implications for infection control. PLoS One. 2011;6(11):e27932. doi: 10.1371/journal.pone.0027932.

12. Mukherjee DV, Cohen B, Bovino ME, Desai S, Whittier S, Larson EL. Survival of influenza virus on hands and fomites in community and laboratory settings. Am J Infect Control. 2012;40(7):590–4. doi: 10.1016/j.ajic.2011.09.006.

13. Bean B, Moore BM, Sterner B, Peterson LR, Gerding DN, Balfour HH, Jr. Survival of influenza viruses on environmental surfaces. J Infect Dis. 1982;146(1):47–51.

14. Shaman J, Kohn M. Absolute humidity modulates influenza survival, transmission, and seasonality. Proc Natl Acad Sci U S A. 2009;106(9):3243–8. doi: 10.1073/pnas.0806852106.

15. Sterling E, Arundel A, Sterling T. Criteria for human exposure to humidity in occupied buildings. ASHRAE transactions. 1985;91(1B):611–22.

16. Koep TH, Enders FT, Pierret C, Ekker SC, Krageschmidt D, Neff KL, et al. Predictors of indoor absolute humidity and estimated effects on influenza virus survival in grade schools. BMC Infect Dis. 2013;13:71. doi: 10.1186/1471-2334-13-71.

17. Lindsley WG, Blachere FM, Davis KA, Pearce TA, Fisher MA, Khakoo R, et al. Distribution of airborne influenza virus and respiratory syncytial virus in an urgent care medical clinic. Clin Infect Dis. 2010;50(5):693–8. doi: 10.1086/650457.

18. Yang W, Elankumaran S, Marr LC. Concentrations and size distributions of airborne influenza A viruses measured indoors at a health centre, a day-care centre and on aeroplanes. J R Soc Interface. 2011;8(61):1176–84. doi: 10.1098/rsif.2010.0686.

19. Minnesota Department of Health Weekly Influenza & Respiratory Illness Activity Report- Summary of 2015-16 Influenza Season. 2016. [updated October 6, 2016; cited 2016 October 31]. Available from: http://www.health.state.mn.us/divs/idepc/diseases/flu/stats/2015summary.pdf.

20. NOAA/NCEP/EMC. North American Land Data Assimilation System. 2016. [cited 2016]. Available from: http://www.emc.ncep.noaa.gov/mmb/nldas.

21. McCoy MH, Wang E. Use of electric cell-substrate impedance sensing as a tool for quantifying cytopathic effect in influenza A virus infected MDCK cells in real-time. J Virol Methods. 2005;130(1-2):157–61. doi: 10.1016/j.jviromet.2005.06.023.

22. Cox CS. Airborne bacteria and viruses. Sci Prog. 1989;73(292 Pt 4):469–99.

23. Tang JW. The effect of environmental parameters on the survival of airborne infectious agents. J R Soc Interface. 2009;6 Suppl 6:S737–46. doi: 10.1098/rsif.2009.0227.focus.

24. Weber TP, Stilianakis NI. Inactivation of influenza A viruses in the environment and modes of transmission: a critical review. J Infect. 2008;57(5):361–73. doi: 10.1016/j.jinf.2008.08.013.

25. Lowen AC, Steel J. Roles of humidity and temperature in shaping influenza seasonality. J Virol. 2014;88(14):7692–5. doi: 10.1128/JVI.03544-13.

26. Yang W, Elankumaran S, Marr LC. Relationship between humidity and influenza A viability in droplets and implications for influenza’s seasonality. PLoS One. 2012;7(10):e46789. doi: 10.1371/journal.pone.0046789.

27. Noti JD, Lindsley WG, Blachere FM, Cao G, Kashon ML, Thewlis RE, et al. Detection of infectious influenza virus in cough aerosols generated in a simulated patient examination room. Clin Infect Dis. 2012;54(11):1569–77. doi: 10.1093/cid/cis237.

28. Blachere FM, Lindsley WG, Slaven JE, Green BJ, Anderson SE, Chen BT, et al. Bioaerosol sampling for the detection of aerosolized influenza virus. Influenza Other Respir Viruses. 2007;1(3):113–20. doi: 10.1111/j.1750-2659.2007.00020.x.

29. Lindsley WG, Schmechel D, Chen BT. A two-stage cyclone using microcentrifuge tubes for personal bioaerosol sampling. Journal of Environmental Monitoring. 2006;8(11):1136–42.

30. Eisfeld AJ, Neumann G, Kawaoka Y. Influenza A virus isolation, culture and identification. Nat Protoc. 2014;9(11):2663–81. doi: 10.1038/nprot.2014.180.

31. Shaman J, Pitzer VE, Viboud C, Grenfell BT, Lipsitch M. Absolute humidity and the seasonal onset of influenza in the continental United States. PLoS Biol. 2010;8(2):e1000316. doi: 10.1371/journal.pbio.1000316.

