## Supplementary Materials for "Humidity as a non-pharmaceutical intervention for influenza A"

### S1 Fig. Methodology flow chart of samples

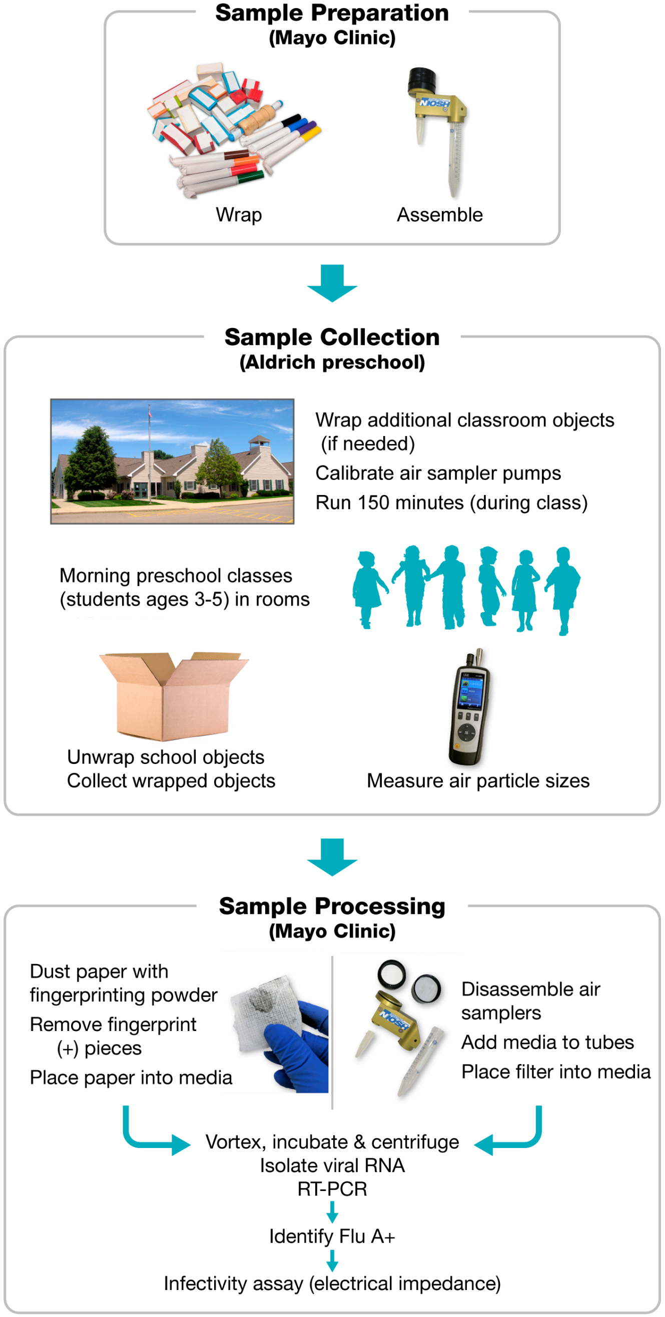

### S2 Fig. Percentage of samples influenza A positive by day by qRT-PCR.

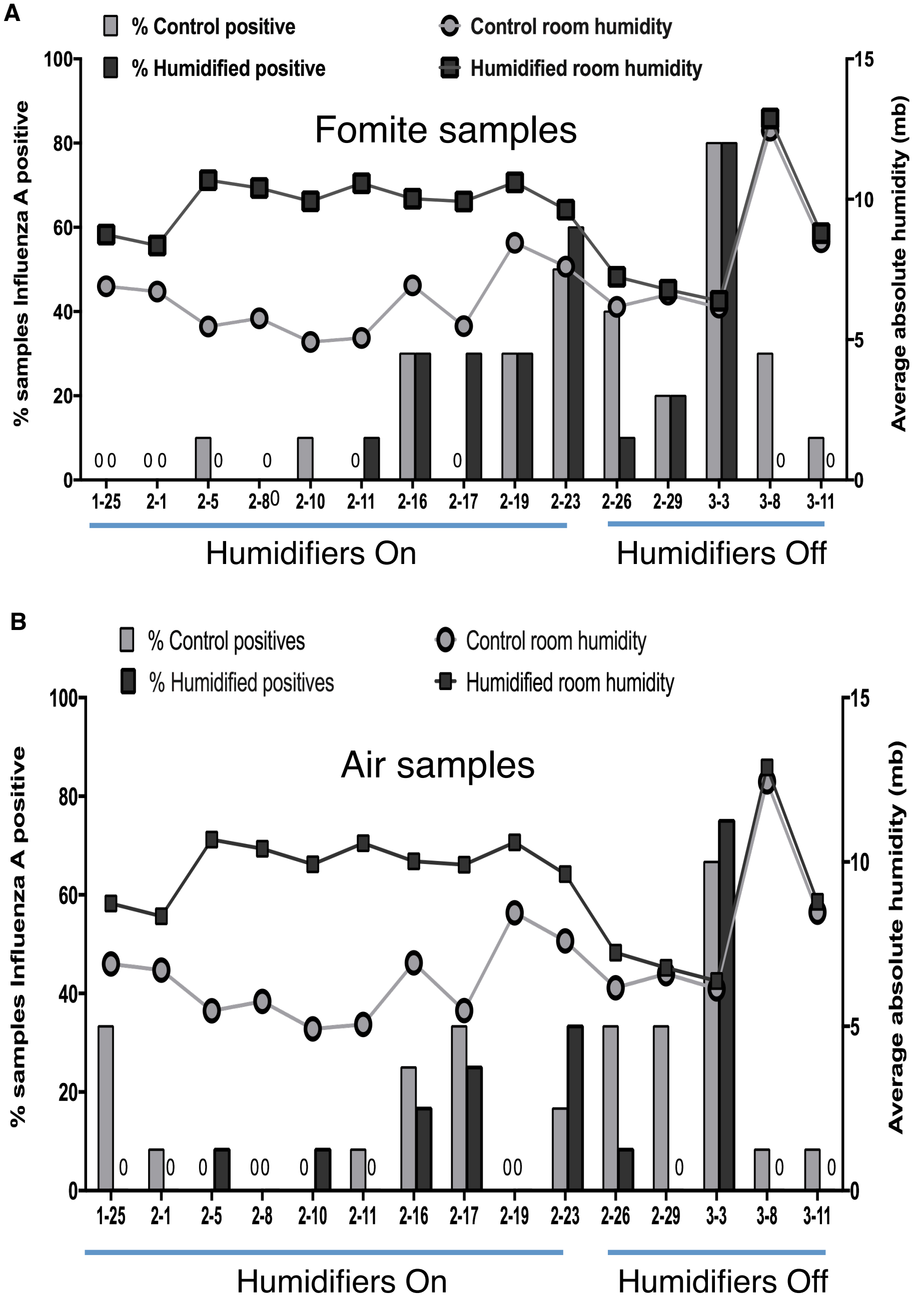


The two lines represent humidity with average of control rooms (grey) and average of humidified rooms (black). The bars are the % of samples positive for influenza A (PCR). (A) Fomite samples (bars), n=10 for control (grey) and humidified (black) rooms except for * where n=5 for control. (B) Air samples (bars), n =24 for control (grey) and humidified (black) rooms.

**S1 Table. Influenza A mean copy number (NS1) by qRT-PCR.**

| Sample | Control [95% CI] | Humidified [95% CI] | P> \|z\| |
|---|---|---|---|
| Fomites | 34.5 [31.96 - 37.11] | 24.6 [22.06 – 27.21] | p<0.001 |
| Air (total) | 41.5 [34.44 – 48.62] | 18.9 [11.84 – 26.02] | p<0.001 |
| Air <1 µm | 8.5 [6.63 - 10.37] | 5.0 [3.13 – 6.87] | p=0.010 |
| Air 1-4 µm | 20.5 [7.73 – 33.34] | 8.9 [-3.90 – 21.70] | p=0.208 |
| Air >4 µm | 12.5 [8.44 – 16.56] | 5.0 [0.98 – 9.09] | p=0.011 |

**S3 Fig. Infectious samples from preschool classrooms during humidification (through 2/23/2016) by electrical impedance assay.**

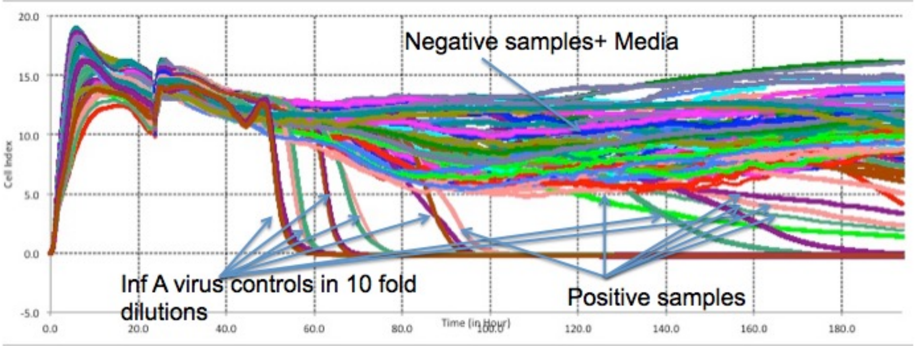


Each line indicates a sample (well) for infectivity. MDCK cells were added at time 0 hours and media changed to samples (n=19 in duplicate), influenza A positive control dilutions (6) in duplicate or control media (n=11) at 24 hours. Additional controls in sample processing (media, paper, fingerprinted paper, hood control) were also included in duplicate as were 12 samples that were PCR negative. Cell indices that returned to 0 indicated cell death (infectious). Arrows indicate individual samples.

**S4 Fig. Floor plans of preschool classrooms including study classrooms.**

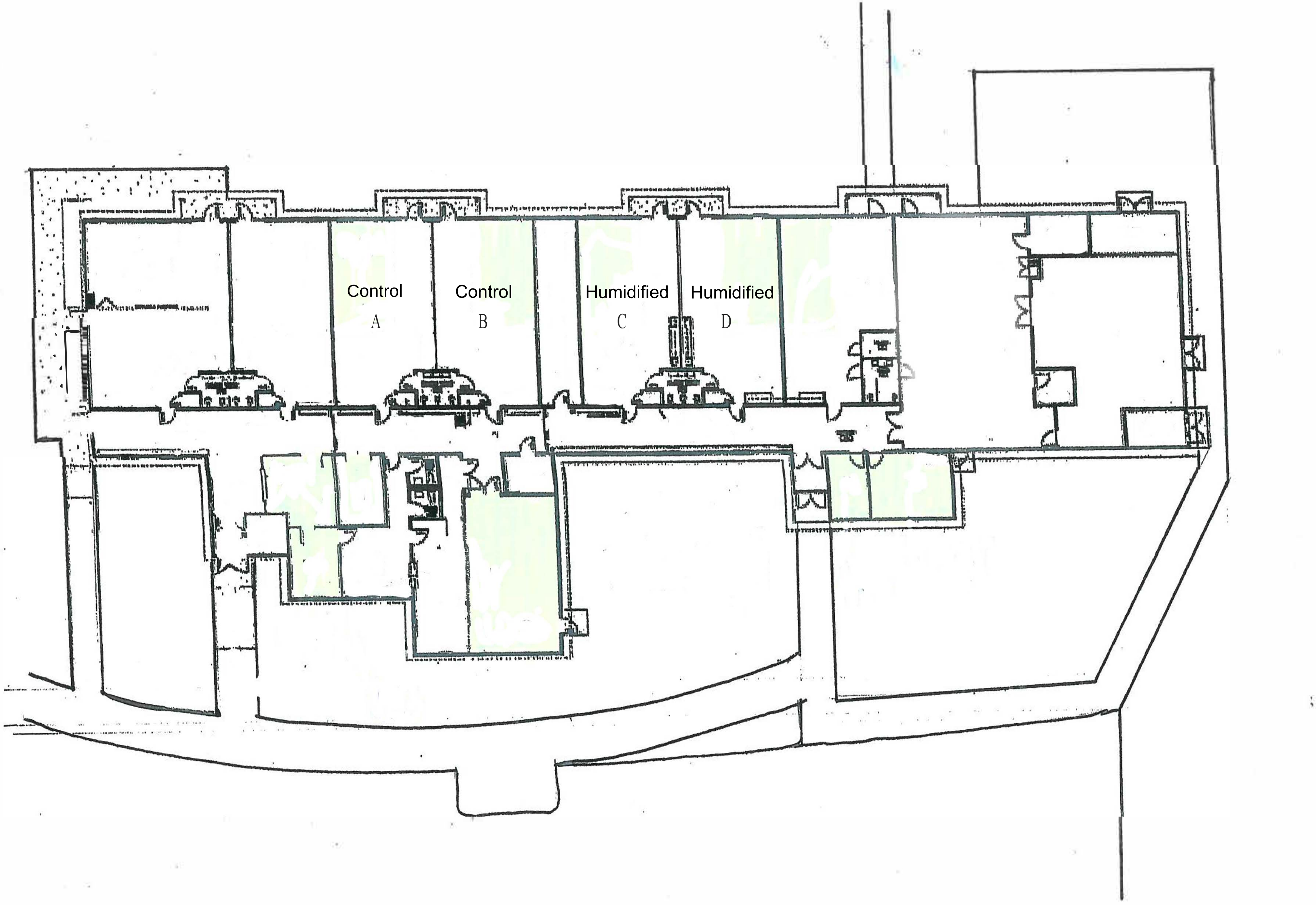

**S2 Table. Sequences of primers used for real-time PCR.**

| Primer name | Primer Sequence | Product size (bp) | Annealing Temperature |
|---|---|---|---|
| InfA F | TTTCACCGAGGAGGGAGCA | 105 | 56.5°C |
| InfA R | CCTCCGATGAGGACCCCAA | | |
| Influenza Bs | GTCCATCAAGCTCCAGTTTT | 145 | |
| Influenza Bas | TCTTCTTACAGCTTGCTTGC | | |
| RSVB F | GCATTAGCCAAAGCAGCAATAC | 155 | |
| RSVB R | CATCCATTAGACCATTTAATTGAGAGC | | |

### S5 Fig. Detection of viral RNA using qRT-PCR.

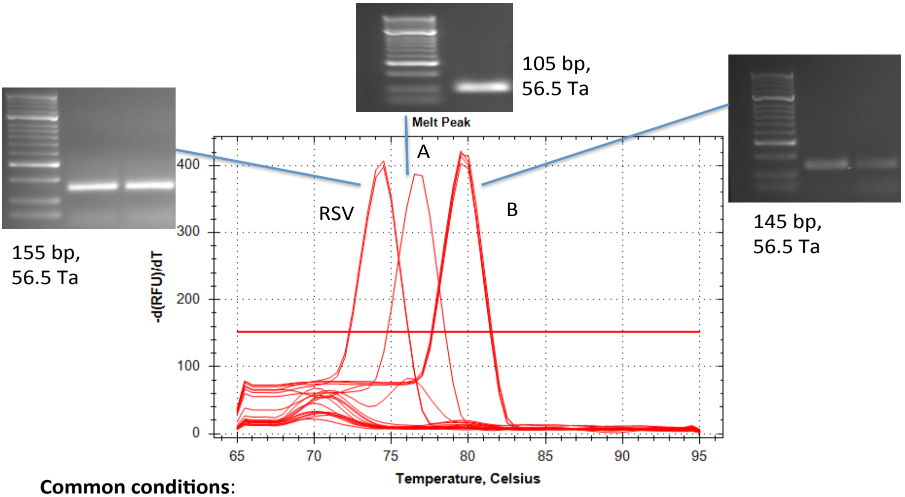


**Common conditions**:

1. Same Ta for all primers
2. Amplicon sizes < 200 bp
3. Tm of each of 3 products differ by 2-3 C

A indicates Influenza A. B indicates Influenza B. RSV indicates respiratory syncytial virus. Ta is annealing temperataure, Tm is melting temperature and bp indicates size in base pairs.

### S6 Fig. Quantification of influenza A positive samples with standard curve.

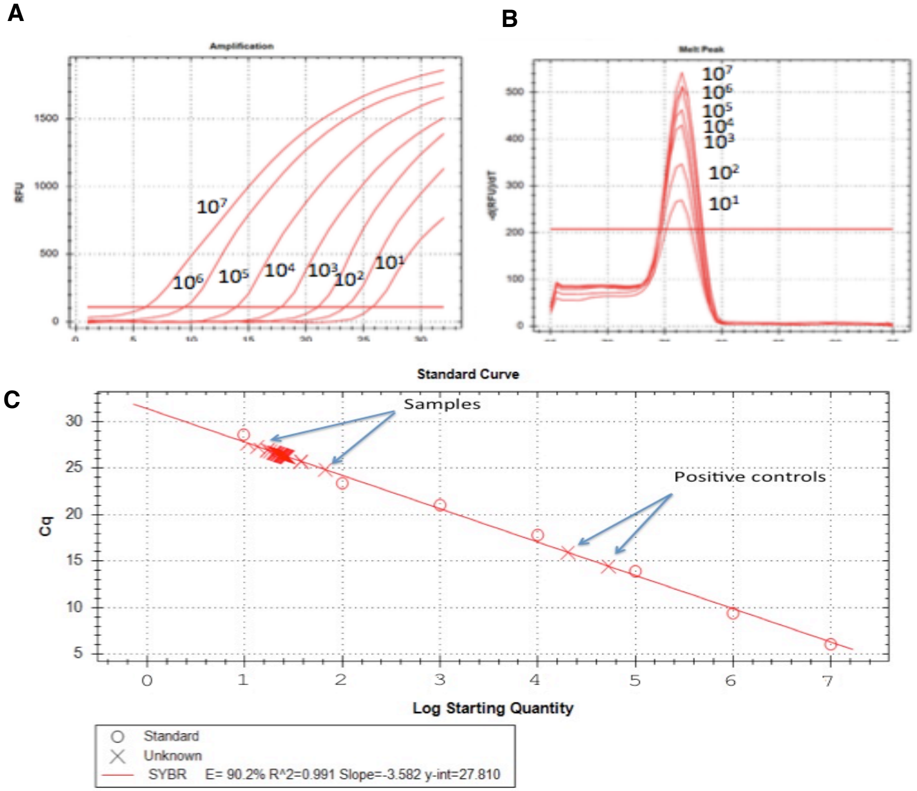


A) Amplification curves of known copy numbers of NS1 gene of infuenza A. B) Melting curves of products from A. C. Standard curve showing standards (O) and experimental samples and positive controls (x) as labeled.
